# A multimodal single-cell atlas of the adolescent brain reveals gene regulatory networks linking development to disease risk

**DOI:** 10.64898/2026.02.20.707029

**Authors:** Isabella C. Galvão, Manuela Lemoine, Tomás V. Waichman, Bhavyaa Chandarana, Steven Hébert, Thais C. de Oliveira, Luciano H. B. dos Santos, Fabio Rogerio, Clarissa L. Yasuda, Enrico Ghizoni, Fernando Cendes, Iscia Lopes-Cendes, Claudia L. Kleinman, Diogo F. T. Veiga

**Affiliations:** Department of Medical Genetics and Genomic Medicine, School of Medical Sciences, Universidade Estadual de Campinas (UNICAMP), Campinas, SP, Brazil; Department of Human Genetics, McGill University, Montreal, Quebec, Canada; Lady Davis Research Institute, Jewish General Hospital, Montreal, Quebec, Canada; Department of Pathology, School of Medical Sciences, Universidade Estadual de Campinas (UNICAMP), Campinas, SP, Brazil; Department of Neurology, School of Medical Sciences, Universidade Estadual de Campinas (UNICAMP), Campinas, SP, Brazil

## Abstract

Adolescence represents a critical window of brain maturation when many neuropsychiatric disorders first emerge, yet the molecular mechanisms driving this developmental period remain incompletely understood. To address this gap, we generated a high-resolution multimodal cell atlas of the developing adolescent brain using paired single-nucleus RNA and ATAC sequencing (snRNA-seq + snATAC-seq) from cortex, hippocampus, and amygdala tissue of six donors aged 6-15 years, profiling 88,658 high-quality nuclei. Integrative analyses identified 36 enhancer-driven gene regulatory networks (eGRNs) with significant age-dependent dynamics in the transition from childhood to adolescence. The majority of adolescence-associated eGRNs were active in oligodendrocytes and their precursors, reflecting active oligodendrogenesis and myelin remodeling during this developmental period. Notably, age-associated cis-regulatory elements were enriched for expression quantitative trait loci (eQTLs) and colocalized with genetic variants linked to both neurodevelopmental and neurodegenerative disorders, suggesting that regulatory networks may be shared across normal adolescent brain development and disease vulnerability. This multimodal cell atlas provides a valuable resource for understanding the human adolescent brain and offers new insights into the molecular origins of neuropsychiatric disorders.

## INTRODUCTION

The transition from childhood to adolescence is characterized by major physical and cognitive changes, including profound neurobiological reorganization in the human brain. Neuroimaging studies have shown that during this developmental period, the human brain undergoes a broad reconfiguration of its cortical macrostructure^1^ and functional connectivity^2^. In this age, the brain circuitry undergoes developmental pruning, a process in which excess axons, dendrites, and synapses generated during early life are selectively eliminated^3^. In parallel, remaining synapses are maintained and strengthened to create an efficient and specialized architecture of mature neural circuits^4^. This refinement is mainly accomplished by glial cells, particularly microglia and astrocytes, with emerging evidence indicating that oligodendrocyte precursor cells also play a role in synaptic remodeling^5^. Additionally, changes in myelin, a lipid bilayer produced by mature oligodendrocytes to insulate axons, have been shown to modulate neuronal networks^6^.

The same plasticity that allows reshaping of neural circuits also carries inherent disease vulnerability. In fact, the majority of psychiatric disorders emerge during this period, with up to 75% arising before 24 years of age^7^. Schizophrenia, for example, most often manifests in adolescence and has been linked to abnormal glia-mediated synaptic pruning and synapse dysfunction^8^.

Despite the importance of this neurodevelopmental period, the cellular and molecular mechanisms driving the transition from the childhood to the adolescent brain remain underexplored. Previous single-cell sequencing studies have generated comprehensive atlases of human neurodevelopment, characterizing early progenitor differentiation, neurogenesis, and gliogenesis during prenatal brain development^9–15^. While these efforts have elucidated the transcriptional and regulatory mechanisms governing fetal cell type diversity and fate specification, postnatal developmental stages remain less well characterized. Notably, a comprehensive single-cell characterization of the human brain focusing on the transition from childhood to adolescence is lacking. This gap leaves fundamental questions unanswered regarding how neural and glial cells continue to mature after birth and how these postnatal developmental processes contribute to the increased vulnerability to neuropsychiatric disorders.

In this study, we applied paired single-nucleus RNA and ATAC sequencing to generate a multimodal cell atlas of the developing adolescent brain to gain insights into the cellular composition and molecular changes associated with human brain development in the period spanning childhood to early adolescence.

As a result, we uncovered age-dependent cell-type-specific regulatory networks that shape cortical and subcortical maturation. Many of these networks involve cis-regulatory regions containing genetic risk variants for neurodevelopmental and neurodegenerative disorders, linking adolescent brain maturation and susceptibility to disease. This cell atlas provides a valuable resource for understanding adolescent brain development at cellular resolution and provides new insights into the molecular foundations of neuropsychiatric disorders.

## RESULTS

### High-resolution multimodal characterization of the developing adolescent brain

We performed paired single-nucleus RNA and ATAC sequencing (snRNA-seq + snATAC-seq) of the amygdala, cortex, and hippocampus from six donors aged 6-15 years, spanning the transition period from childhood to early adolescence, to characterize the cellular composition and molecular changes in the developing adolescent human brain (Figure 1A, Table 1). After strict quality control, a total of 88,658 nuclei with paired transcriptomic and chromatin accessibility profiles were retained for downstream analyses (Figure 1B). Genetic ancestry analysis based on whole-genome sequencing indicated that our pediatric donors were admixed, showing a predominant European genetic background with African and Indigenous American contributions, as previously reported for the Brazilian population^16^ (Methods, Figure S1A-D).

**Figure 1.**
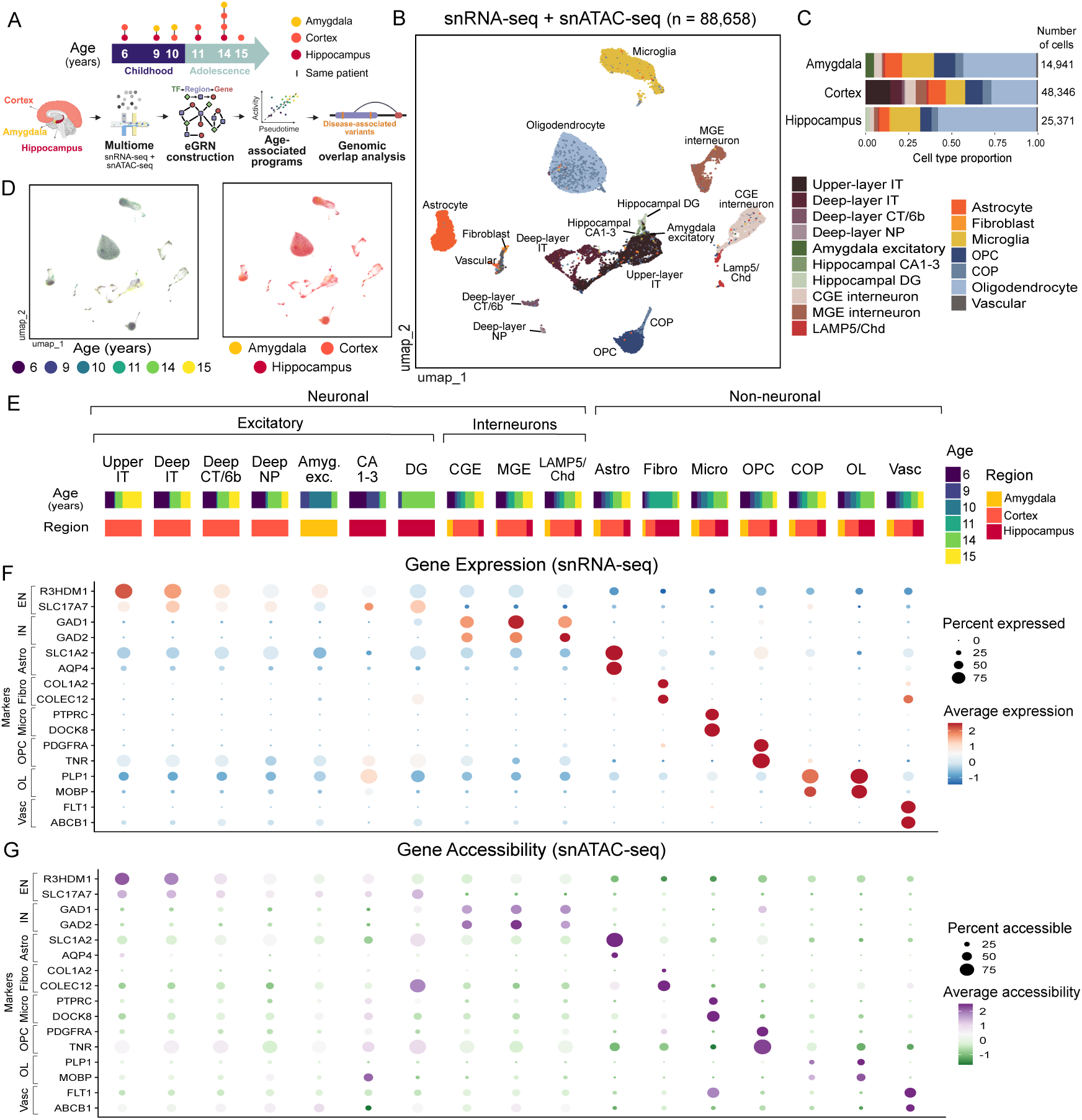
Multimodal single-nucleus profiling of the developing adolescent brain. (A) Overview of study design. Nuclei were isolated from the amygdala, cortex, and hippocampus of six pediatric donors (ages 6–15 years) and profiled using paired single-nucleus RNA sequencing (snRNA-seq) and single-nucleus assay for transposase-accessible chromatin sequencing (snATAC-seq). Integrated data analysis enabled the identification of enhancer-driven gene regulatory networks and age-associated cellular programs. Genomic overlap analysis of these programs revealed enrichment for disease-associated variants. Schematic created with BioRender. (B) Joint Uniform Manifold Approximation and Projection (UMAP) embedding of snRNA-seq and snATAC-seq data, colored by annotated cell type. (C) Barplots denoting cell type distribution in the amygdala, cortex, and hippocampus. (D) UMAP embedding of snRNA-seq and snATAC-seq data, colored by age (left) and brain region (right). (E) Barplots denoting cell type distribution according to age (left) and brain region (right). (F) Dot plot of gene expression (scRNA-seq) for canonical markers of major cell types. Dot size represents the fraction of cells expressing the marker, and color indicates the average expression level. (G) Dot plot of chromatin accessibility (scATAC-seq) for cell-type marker genes shown in (F). Dot size denotes the fraction of cells with open chromatin for the marker gene, and color represents the average chromatin accessibility. Upper IT, upper-layer intratelencephalic; Deep IT, deep-layer intratelencephalic; Deep CT/6b, deep-layer corticothalamic/6b; Deep NP, deep-layer near-projecting; Amyg. exc, amygdala excitatory; CA1–3, hippocampal CA 1–3; DG, hippocampal dentate gyrus; CGE, caudal ganglionic eminence interneuron; MGE, medial ganglionic eminence interneuron; LAMP5/Chd, LAMP5+ chandelier interneuron; Astro, astrocyte; Fibro, fibroblast; Micro, microglia; OPC, oligodendrocyte precursor cell; COP, committed oligodendrocyte precursor cell; OL, oligodendrocyte; Vasc, vascular cells.

**Table 1.**
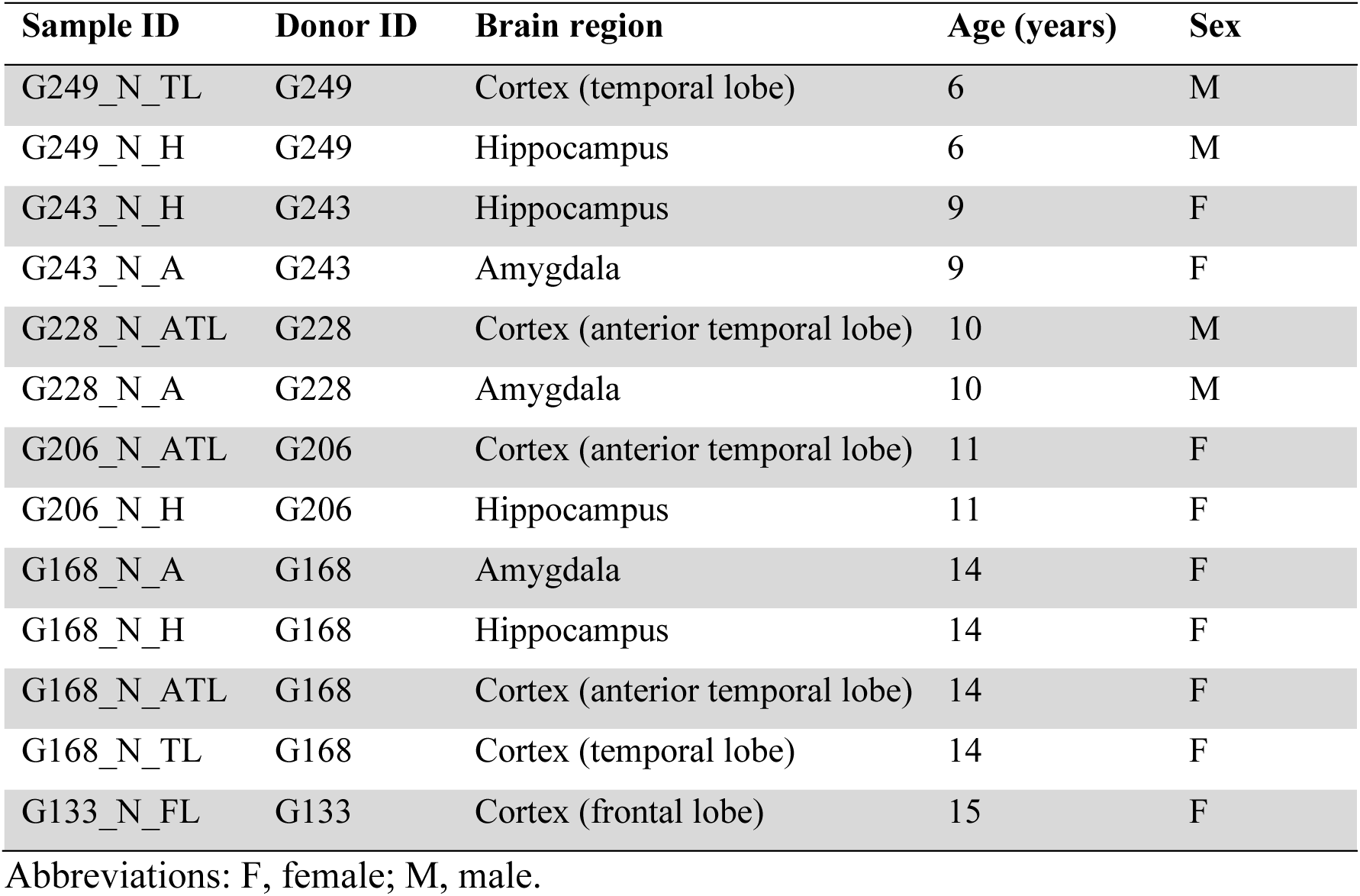
Demographic and clinical characteristics of brain tissue samples used for single-cell multiome profiling.

| Sample ID | Donor ID | Brain region | Age (years) | Sex |
| --- | --- | --- | --- | --- |
| G249_N_TL | G249 | Cortex (temporal lobe) | 6 | M |
| G249_N_H | G249 | Hippocampus | 6 | M |
| G243_N_H | G243 | Hippocampus | 9 | F |
| G243_N_A | G243 | Amygdala | 9 | F |
| G228_N_ATL | G228 | Cortex (anterior temporal lobe) | 10 | M |
| G228_N_A | G228 | Amygdala | 10 | M |
| G206_N_ATL | G206 | Cortex (anterior temporal lobe) | 11 | F |
| G206_N_H | G206 | Hippocampus | 11 | F |
| G168_N_A | G168 | Amygdala | 14 | F |
| G168_N_H | G168 | Hippocampus | 14 | F |
| G168_N_ATL | G168 | Cortex (anterior temporal lobe) | 14 | F |
| G168_N_TL | G168 | Cortex (temporal lobe) | 14 | F |
| G133_N_FL | G133 | Cortex (frontal lobe) | 15 | F |

Abbreviations: F, female; M, male.

Cell type annotation using an ensemble of prediction models trained on a reference dataset^17^ containing brain cell types from the cortex, amygdala, and hippocampus identified 17 distinct cell types (Methods, Figure 1B). Most of the sequenced cells were derived from the cortex, followed by the hippocampus and amygdala (Figure 1C). Excitatory neurons were mainly region-specific and comprised cortical subpopulations such as upper/deep-layer intratelencephalic (IT), deep-layer near-projecting (NP), deep-layer corticothalamic (CT/6b), as well as hippocampal-specific subpopulations such as CA1–3 and dentate gyrus (DG) and amygdala excitatory neurons. Interneurons (i.e., inhibitory neurons) derived from the amygdala, cortex, and hippocampus, and included caudal ganglionic eminence (CGE), medial ganglionic eminence (MGE), and LAMP5⁺/Chandelier subtypes. Non-neuronal populations included astrocytes, fibroblasts, microglia, oligodendrocyte precursor cells (OPCs), committed oligodendrocyte precursors (COPs), and mature oligodendrocytes. Clustering analysis revealed 42 clusters with distinct transcriptional and epigenetic profiles (Figure S2A), each predominantly composed of a unique consensus cell type, validating the single-cell level annotation (Figure S2B). Additionally, no cluster or cell type was specific to a single donor, brain region (aside from region-specific excitatory neurons), or biological age (Figures 1D, S2C-D).

To confirm cell type identity, we also examined the gene expression and chromatin accessibility of canonical marker genes for major cell types (Figures 1F–G). Gene expression patterns of canonical markers were mostly concordant with chromatin accessibility, indicating coordinated regulation at the transcriptomic and epigenomic levels. Of note, canonical markers of excitatory neurons were more highly expressed in cortical neurons than in their hippocampal and amygdala counterparts.

The cellular composition was notably distinct in the amygdala, cortex, and hippocampus, indicating region-specific distributions of brain cell types (Figures 1E, S3A). The neuronal density was highest in the cortex (∼37% compared to approximately 10% in both the amygdala and hippocampus). Excitatory neuron subtypes exhibited regional specificity, and among interneuron populations, CGE-derived interneurons were particularly enriched in the amygdala. Astrocytes were most prevalent in the amygdala, followed by the cortex and hippocampus. Fibroblasts were primarily detected in the hippocampus, although with substantial donor-to-donor variability. Microglia were more abundant in the amygdala and hippocampus (∼20%) compared to the cortex (∼10%). In the oligodendrocyte lineage, oligodendrocyte precursor cells (OPCs) were more frequent in the amygdala. In contrast, mature oligodendrocytes were more abundant in the hippocampus than in either the amygdala or the cortex, suggesting that OPC differentiation may occur earlier in the hippocampus. Thus, aside from region-specific excitatory neurons, all cell types were detected with varying frequencies across the analyzed brain regions.

### Cellular composition is maintained in the transition from childhood to adolescence

Next, we aimed to compare the cellular composition between childhood and adolescent brains across different regions. We classified cells based on their developmental stage into childhood (≤10 years) and adolescence (>10 years), and performed statistical testing to compare cell type frequency between these stages. Because brain regions are major determinants of cellular composition, all statistical tests were performed separately for the cortex, hippocampus, and amygdala. Overall, we found no significant changes in cell type frequency (Figure S3B), indicating no major shifts in the brain cellular landscape during the transition from childhood to adolescence.

### eGRNs mapping identifies core transcriptional regulators in brain cell types

To study the transcriptional regulation during the shift from childhood to adolescence in the human brain, we used SCENIC+ to combine chromatin accessibility with gene expression data to identify enhancer-driven gene regulatory networks (eGRNs). SCENIC+ finds regulatory triplets—combinations of transcription factors (TFs), their related cis-regulatory elements (enhancers/open chromatin regions), and downstream target genes—mapping the regulatory landscape within different brain cell types. The group of cis-regulatory elements and their downstream target genes regulated by a given TF is called an eRegulon.

The UMAP embedding based on eRegulons activity scores grouped cells primarily by cell type, indicating that TF activity is a key determinant of cellular identity (Figure 2A). Notably, SCENIC+ uncovered known master regulators of neuronal and glial populations. For instance, *CUX2* and *FOXP1* eRegulons were found to be specifically activated in excitatory neurons, *DLX1* and *LHX6* in interneurons, *SOX9* and *TCF7L1* in astrocytes, *IRF8* and *CEBPD* in microglia, *OLIG2* and *ETV5* in OPCs, and *SOX10* and *OLIG1* in oligodendrocytes (Figures 2B,C).

**Figure 2.**
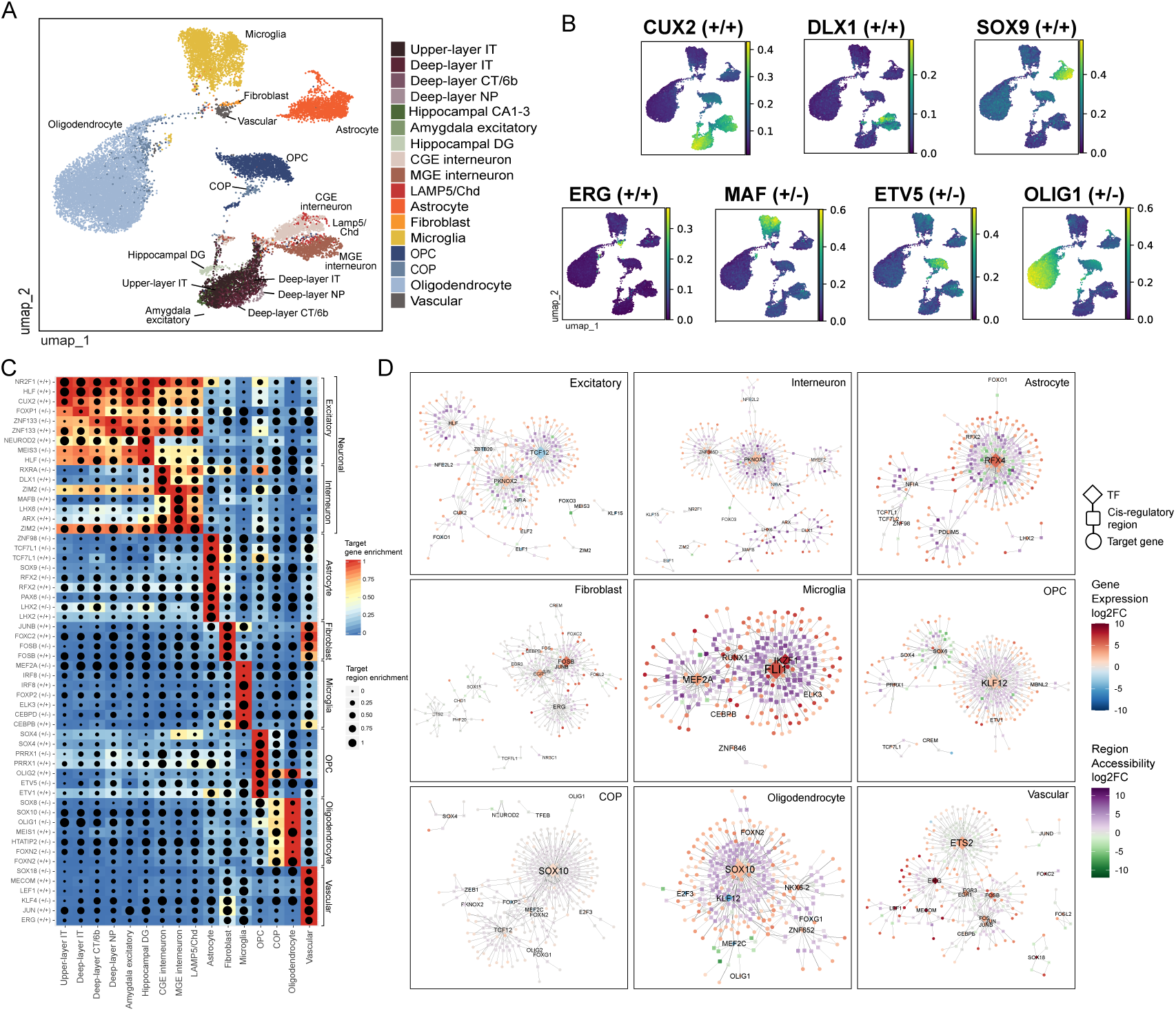
Cell–type–resolved enhancer-driven gene regulatory networks (eGRNs) in the adolescent brain. (A) UMAP visualization of nuclei based on eRegulons activity (AUC score), colored by annotated cell type. (B) Feature plots showing eRegulons activity (AUC score) of established master transcriptional regulators of brain cell types. eRegulon names are defined by the driving transcription factor followed by the sign of the correlation with the target genes and target regions, respectively. A plus sign (+) indicates a positive correlation and a minus sign (−) indicates a negative correlation. (C) Heatmap displaying cell type–specific eRegulons, defined as those with the highest Regulon Specificity Scores (RSS) for each cell type. RSS quantifies the degree to which an eRegulon is specifically active within a given cell type. Tile color represents target gene enrichment (gene-based AUC), and dot size reflects target region enrichment (region-based AUC). (D) Network visualization of eGRNs reconstructed for brain cell types. The top-ranked 200 regulatory interactions - triplets of TF, cis-regulatory region, and target gene-were selected among the 20 most specific eRegulons (highest RSS) per cell type. TFs are depicted as diamonds, cis-regulatory regions as squares, and target genes as circles. Node colors indicate log₂ fold-changes in chromatin accessibility (regions) and gene expression (genes). OPC, oligodendrocyte precursor cell; COP, committed oligodendrocyte precursor cell.

In addition, network analysis of the top eRegulons revealed regulatory hubs across brain cell types, uncovering master regulators and cooperation among core TFs (Figure 2D). For instance, PKNOX2 emerged as a master regulator in both excitatory neurons and interneurons, consistent with its established role in neuronal differentiation^18^. RFX4 was identified as a master regulator in astrocytes, although its functions in postnatal astrocytes remain poorly understood. FLI1 and IKZF1, which are known to regulate cytokine production during inflammation and modulate synaptic pruning through inflammasome activity^19,20^, were identified as core regulators in microglia. In OPCs, KLF12 emerged as the key regulator, consistent with its role in promoting OPC differentiation and supporting white-matter repair^21^. SOX10, which regulates terminal differentiation and myelination^22^, was identified as a core regulator in both COPs and mature oligodendrocytes.

### Pseudotime analysis of eGRNs activity uncovers regulatory networks associated with the developing adolescent brain

To identify regulatory networks associated with adolescent brain development, we analyzed eRegulon activity along a pseudotime trajectory spanning childhood to adolescence and identified eRegulons with significant activity during this transition (see Methods). We focused on activator eRegulons (those in which TF expression leads to increased target gene expression) whose activity showed a significant linear correlation with age. This analysis revealed 36 eRegulons exhibiting significant age-dependent dynamics during the transition from childhood to adolescence, in both neuronal and glial cell types (Figure 3, Supplementary Tables 1,2).

**Figure 3.**
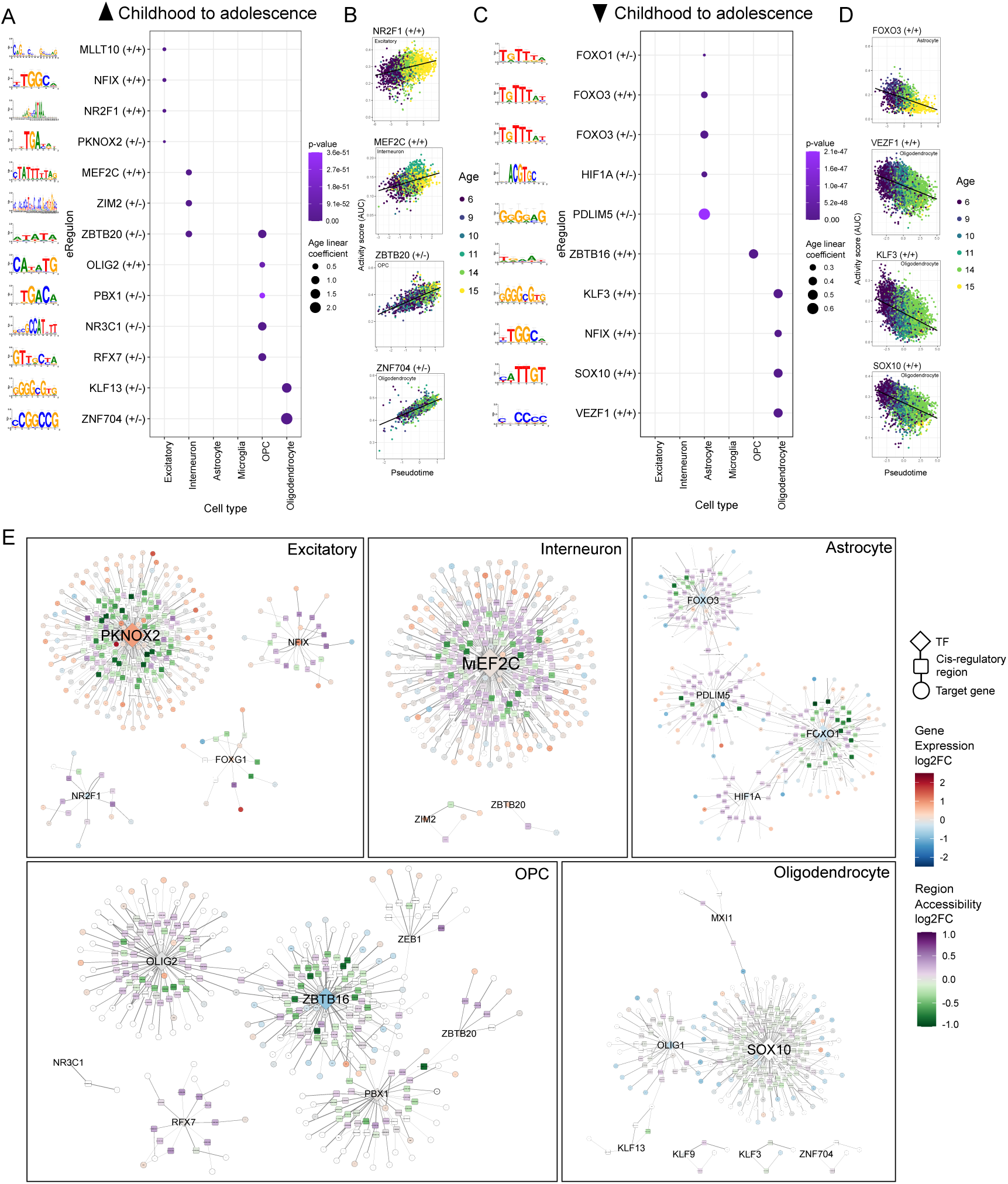
Age-dependent transcriptional regulation in the adolescent brain. (A–B) eRegulons predicted to have increased activity during the transition from childhood to adolescence. (A) Dot plot showing eRegulons with significant age-dependent dynamics. Dot size reflects the age coefficient, and color indicates the p-value from a linear model of z-scored eRegulon activity versus pseudotime. Sequence motifs recognized by transcription factors (TFs) in the eRegulon are shown on the left. eRegulon names are defined by the driving TF followed by the sign of the correlation with the target genes and target regions, respectively. A plus sign (+) indicates a positive correlation and a minus sign (−) indicates a negative correlation. (B) Representative scatter plots of eRegulon activity (AUC) along pseudotime for selected cell types. Point color represents donor age, and the black line denotes the fitted linear trend. (C-D) eRegulons predicted to have decreased activity during the transition from childhood to adolescence. Legends are defined as in (A) and (B), respectively. (E) Network visualization of age-dependent eGRNs for brain cell types. The top-ranked 200 regulatory interactions - triplets of TF, cis-regulatory region, and target gene - were selected among age-associated eRegulons per cell type. TFs are depicted as diamonds, cis-regulatory regions as squares, and target genes as circles. Node colors indicate log₂ fold-changes in chromatin accessibility (regions) and gene expression (genes).

Among these, 16 eRegulons showed increased activity (top eRegulons shown in Figures 3A,B) and 20 showed decreased activity with age (top eRegulons shown in Figures 3C,D). Most adolescence-associated eRegulons were active in a particular cell type, with only two —*NFIX* and *ZBTB20*— shared across multiple lineages. Notably, the majority of adolescence-associated eRegulons were detected in oligodendrocytes (16/36) and OPCs (7/36), consistent with ongoing oligodendrogenesis and myelin remodeling known to occur during this developmental window. For instance, KLF13, which is specifically expressed in oligodendrocytes (Figure S4), has been shown to control the expression of myelin-associated genes by binding to regulatory regions also recognized by SOX10^23^, suggesting coordinated activity between these eRegulons. Additional adolescence-associated eRegulons were detected in astrocytes (5/36), excitatory neurons (5/36), and interneurons (3/36). Notably, astrocytic regulators were mainly FOXO-family TFs, which are known to control astrocyte metabolism^24,25^. Network visualization of top-ranked regulatory triplets (TF-region-genes) reveals some degree of cooperation among age-related TFs, such as the FOXO1/FOXO3/PDLIM5 in astrocytes, and SOX10/OLIG1 in oligodendrocytes (Figure 3E). Altogether, the eRegulon trajectory analysis identified active brain regulatory networks and highlighted oligodendrocyte and OPC transcriptional regulation as major transcriptional events associated with brain maturation during the transition from childhood to early adolescence.

### eGRNs in the developing adolescent brain are enriched in genetic variants associated with neuropsychiatric risk

Next, we asked whether cis-regulatory elements in adolescence-associated eRegulons were known to control gene expression in the brain. We overlapped cis-regulatory regions associated with adolescent brain development with MetaBrain, a comprehensive resource of expression quantitative trait loci (eQTLs) containing 3,549 genetic variants influencing gene expression across multiple brain regions, including cortex, hippocampus, and amygdala.

The cis-regulatory regions in adolescence-associated eRegulons showed a significant enrichment for brain eQTLs (P < 10⁻⁴; Supplementary Table 3). For instance, *S100B*, a gene involved in oligodendrocyte maturation^26,27^ and part of the SOX10, KLF13, and OLIG1 eRegulons activated in oligodendrocytes, is controlled by multiple cis-regulatory elements overlapping proximal and distal ENCODE enhancers, including a region 16 Kb upstream of the promoter containing a known eQTL (Figure 4A,B). Thus, the significant enrichment of eQTLs within regulatory regions of the adolescent brain suggests their functional role in controlling gene expression during this period.

**Figure 4.**
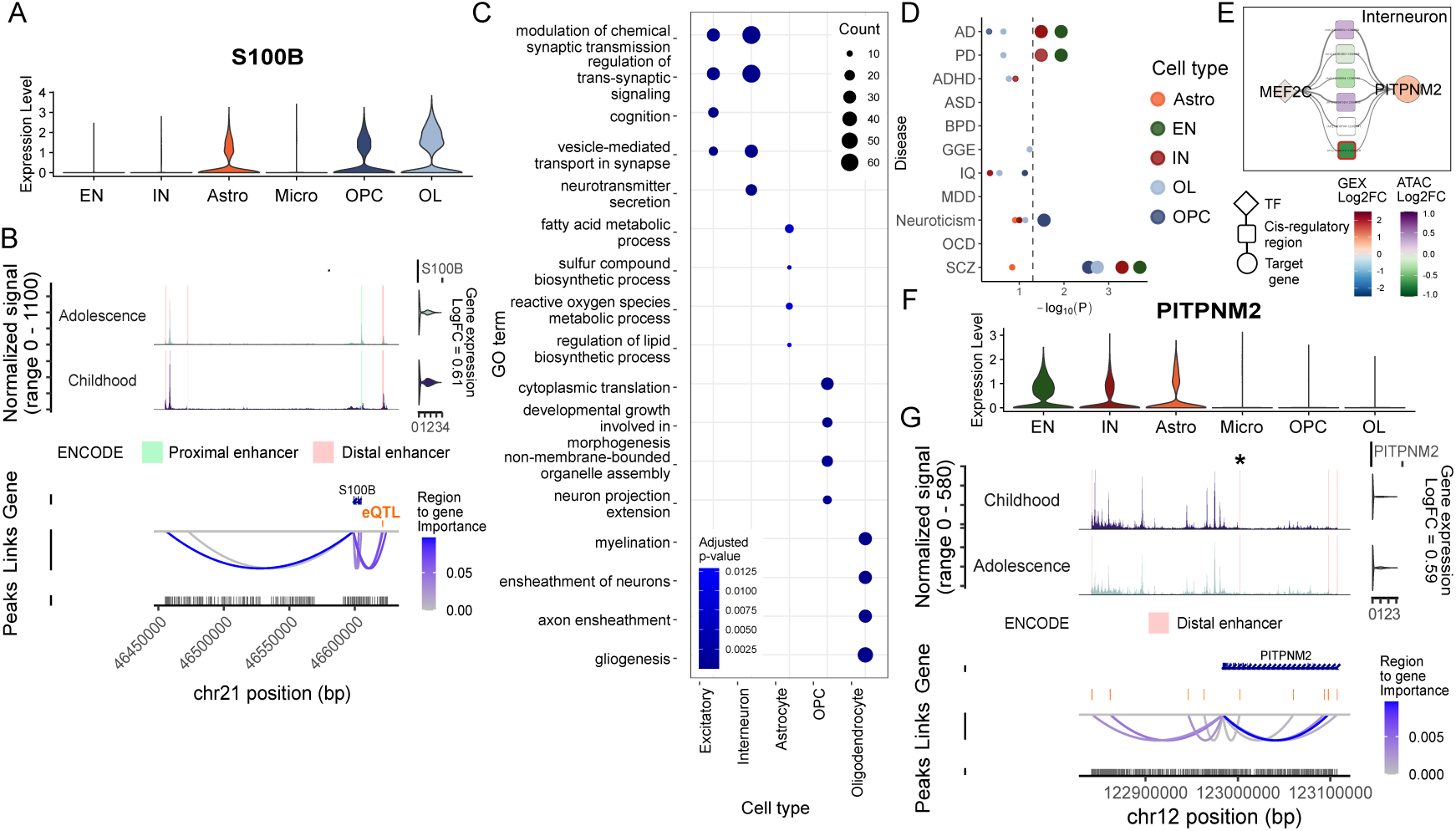
Functional characterization of adolescence-associated eRegulons. (A) Violin plots of *S100B* gene expression across cell types. (B) Regulatory links for *S100B* inferred by SCENIC+ in oligodendrocytes. Chromatin accessibility (snATAC-seq) and expression (scRNA-seq) for *S100B* in childhood (≤ 10 years) and adolescence (> 10 years) are shown at the top. Vertical bars indicate regions annotated as ENCODE enhancers. The gene track displays the genomic location of the *S100B* locus on chromosome 21, and the bottom track shows accessible chromatin peaks within this region. Link color denotes region-to-gene importance scores, and the orange line highlights the overlap with an expression quantitative trait locus (eQTL) associated with *S100B* regulation. (C) Dot plot displaying enriched Gene Ontology (GO) terms for target genes in age-associated eRegulons according to cell type, computed using clusterProfiler. Dot size and color correspond to the number of genes associated with the GO term and enrichment p-value, respectively. (D) Dot plot showing enrichment of disease-associated variants within cis-regulatory regions of adolescence-associated eRegulons. The *x*-axis represents the overlap significance (−log10 of the adjusted *P* value) from permutation-based enrichment tests, and the *y*-axis lists the corresponding diseases. Dot color denotes the cell type in which the enrichment was detected, and the dashed line marks the significance threshold (P < 0.05). (E) Network visualization of *PITPNM2* transcriptional regulation by MEF2C in interneurons. TFs are depicted as diamonds, cis-regulatory regions as squares, and target genes as circles. Node colors indicate log₂ fold-changes in chromatin accessibility (regions) and gene expression (genes). Red borders indicate differentially accessible regions. (F) Violin plots of *PITPNM2* gene expression across cell types. (G) Regulatory links for *PITPNM2* inferred by SCENIC+ in interneurons. Orange lines next to regulatory links denote the overlap with multiple genetic risk variants associated with schizophrenia. See legend in (B). EN, excitatory neuron; IN, interneuron; Astro, astrocyte. Micro, microglia; OPC, oligodendrocyte precursor cell; OL, oligodendrocyte; AD, Alzheimer’s disease; PD, Parkinsons’s disease; ADHD, attention deficit hyperactivity disorder; ASD, autism spectrum disorder; BPD, bipolar disorder; GGE, generalized genetic epilepsy; IQ, intelligence quotient; MDD, major depressive disorder; OCD, obsessive compulsive disorder; SCZ, schizophrenia.

In addition, functional annotation of eRegulons target genes using the Gene Ontology revealed enrichment of key cellular processes in brain cells including modulation of synaptic signaling (excitatory neurons and interneurons), cognition (excitatory neurons), lipid and mitochondrial metabolism (astrocytes), developmental cell growth (OPCs), gliogenesis (OPCs and oligodendrocytes), and myelination (oligodendrocytes) (Figure 4C, Supplementary Table 4).

To explore the potential relevance of these eGRNs to neurological disease, we also overlapped the cis-regulatory regions associated with adolescent brain development with genetic risk variants reported in previous genome-wide association (GWAS) studies (Supplementary Table 5). We observed significant enrichment for variants linked to Alzheimer’s and Parkinson’s disease in adolescence-associated eRegulons from excitatory neurons and interneurons, neuroticism in OPCs, and schizophrenia in excitatory neurons, interneurons, OPCs, and oligodendrocytes (Figure 4D; Supplementary Table 6). For example, we found that *PITPNM2*, a target of the MEF2C eRegulon that has increased activity in adolescence in interneurons, is controlled by multiple cis-regulatory regions containing genetic risk variants associated with schizophrenia (Figures 4E-G). *PITPNM2* expression was detected in excitatory neurons, interneurons, and astrocytes (Figure 4F), with a marked upregulation specifically in interneurons from childhood to adolescence. Notably, we found nine GWAS variants within cis-regulatory regions inferred by SCENIC+, as highlighted by the orange lines next to regulatory links in Figure 4G. Also, several cis-regulatory elements linked to *PITPNM2* were found to be overlapping ENCODE-annotated distal enhancers, including a region that showed differential chromatin accessibility in adolescence, suggesting that this gene is actively regulated by MEF2C during this developmental period (Figure 4G). Thus, these observations suggest that *PITPMN2* is critical for normal brain maturation in adolescence and might be associated with schizophrenia, as this locus is a hotspot of GWAS variants.

Collectively, these results from the brain eQTL and risk variants analyses suggest that cis-regulatory elements in adolescence-associated eGRNs are enriched for genetic variants that influence both normal adolescent brain maturation and disease susceptibility.

## DISCUSSION

In this study, we used multimodal single-nucleus profiling to characterize transcriptional and epigenomic programs underlying the transition from childhood to adolescence across the amygdala, cortex, and hippocampus. By exploring cellular diversity and cell-type-specific regulatory landscapes through enhancer-driven gene regulatory networks, we uncovered cellular programs that modulate adolescent brain maturation during this critical developmental window of the human brain.

Analyses of cellular composition revealed regional differences but no evidence of age-specific cell populations emerging during this period, suggesting that late childhood and adolescence are characterized by regulatory remodeling rather than major cellular shifts. In contrast, previous single-cell studies have reported age-related changes, particularly within the oligodendrocyte lineage, including a decline in OPCs accompanied by an increase in mature oligodendrocytes from infancy into adulthood^28,29^. On the other hand, our study focused on a narrower age period specifically associated with the shift from childhood to adolescence, which might prevent the detection of later cellular changes.

Single-cell multiomics quantifies molecular features including gene expression and chromatin accessibility within individual cells. Joint analysis of these multimodal single-cell measurements enables linking cis-regulatory elements to target genes to elucidate how regulatory networks control gene expression at cell-type resolution. Using this approach, we identified enhancer-driven regulons (eRegulons) and core regulators for each major brain cell type. We found that master regulators are largely conserved across regions and enriched for functions related to cellular differentiation, specialization, survival, and homeostasis^18–23,30–34^. Notably, many of these regulators have established links to neurological disease. For example, RFX4, identified here as a core astrocytic regulator, has been implicated in both neurodevelopmental disorders such as autism spectrum disorder and attention-deficit/hyperactivity disorder^35^, as well as in neurodegenerative conditions including Alzheimer’s disease^36^.

To capture regulatory dynamics specific to adolescent brain development, we identified eRegulons with differential activity in the transition from childhood to adolescence. These adolescence-associated eRegulons were predominantly active in oligodendrocyte lineage cells, thus recapitulating prior studies showing that oligodendrogenesis and myelin remodeling transcriptional programs are active processes in late postnatal development^37^. In addition, adolescence-associated regulatory programs were detected in neurons and astrocytes, where they were enriched for pathways related to synaptic signaling, cognition, and metabolic regulation. Unexpectedly, we did not detect adolescence-associated regulatory programs in microglia, despite their well-established role in synaptic pruning during development, including in early adolescence^38^. This suggests that synaptic pruning mediated by core microglia regulators is maintained continuously from childhood to adolescence, rather than exhibiting an age-dependent shift during this window.

Notably, we found that adolescence-activated eRegulons are strongly enriched for genetic variants associated with schizophrenia, and to a lesser extent with Alzheimer’s disease, and Parkinson’s disease. A prominent example is PITPNM2 gene in the MEF2C regulon, which was found to be upregulated in adolescence in a locus with multiple cis-regulatory elements harboring schizophrenia-associated variants. Moreover, GRIN2A, the first monogenic cause of early-onset neuropsychiatric disorders^39^, was identified within these adolescent-associated programs. These findings suggest that regulatory networks active during normal adolescent brain development may also confer susceptibility to neuropsychiatric and neurodegenerative disorders. In addition, understanding the functional consequences of risk variants on gene regulation remains challenging, and cell-type-specific gene networks learned from single-cell multiomics provide a framework for variant interpretation and prioritization.

Our study has limitations. Our method captures cell-type regulatory programs that show only consistent linear correlation with age, since detecting non-linear dynamics in smaller cohorts might be affected by donor variability. Future studies leveraging larger cohorts are warranted to further refine and expand regulatory networks underlying adolescent brain maturation, including the construction of region-specific brain networks.

Overall, this multimodal cell atlas provides a valuable resource for dissecting the cellular and regulatory landscapes of adolescent brain maturation and for prioritizing follow-up studies of genetic risk variants in neuropsychiatric disease.

## MATERIALS AND METHODS

### Clinical samples and neuropathological evaluation

Brain tissue from the amygdala, cortex, and hippocampus was collected from six donors undergoing epilepsy surgery at the Hospital de Clínicas, University of Campinas. The study was approved by the Institutional Research Ethics Board (CAAE: 12112913.3.0000.5404), and written informed consent was obtained from all children or their legal guardians before surgery. For tissue resected during epilepsy surgery, periepileptic regions were selected for experiments following histopathological examination that confirmed normal cytoarchitecture and the absence of cellular abnormalities.

### Whole-genome sequencing

Genomic DNA was extracted from brain tissue using the PureLink Genomic DNA Mini Kit (Invitrogen, Cat. No. K182001) according to the manufacturer’s instructions. Sequencing libraries were prepared using the TruSeq Nano DNA Library Preparation Kit (Illumina, Part no. 15041110 Rev. D). Paired-end sequencing (151 bp) was performed on the Illumina NovaSeq 6000 platform, targeting 30× genome coverage.

### Genetic ancestry analysis

We used the publicly available Human Genome Diversity Project (HGDP) + 1000 Genomes Project (1KGP) reference panel from gnomAD v3.1.2, which comprises 4,151 individuals from global populations representing major continental groups: Africa, Europe, Central South Asia, East Asia, the Middle East, Oceania, Indigenous American, and admixed American populations^40^. Quality control filtering was performed separately on the reference panel and the Brazilian samples sequenced in this study prior to merging using PLINK v2.0^41^. For the HGDP+1KGP reference panel, we applied the following filters: removal of variants failing quality filters; minor allele frequency (MAF) < 0.01, Hardy-Weinberg equilibrium (HWE) exact test p-value < 10⁻⁸; and restriction to biallelic autosomal SNPs. For the Brazilian samples, we applied similar filters with a less stringent HWE threshold (p < 10⁻⁵) to account for the smaller sample size. After individual-level filtering, we identified the intersection of SNPs present in both datasets. The datasets were then merged using PLINK v1.9^42^, and SNPs with genotype missingness > 5% were removed. To obtain a set of independent SNPs for population structure analyses, we performed linkage disequilibrium (LD) pruning using PLINK v2.0 with a sliding window of 50 SNPs, a step size of 10 SNPs, and an r² threshold of 0.1, yielding a final dataset of 315,983 autosomal SNPs. Principal component analysis (PCA) was performed on the LD-pruned dataset using SNPRelate^43^ to assess population structure and visualize the genetic relationship between the Brazilian samples and global reference populations. Global ancestry proportions were estimated using ADMIXTURE v1.3.0^44^ with the LD-pruned dataset. We performed unsupervised clustering analyses for *K*=2 to *K*=6 ancestral populations using 5-fold cross-validation. The optimal *K* value was determined based on the lowest cross-validation error.

### Nuclei isolation and library preparation

Nuclei were isolated using the Chromium Nuclei Isolation Kit (10X Genomics) following the manufacturer’s protocol (CG000505, Rev. A), compatible with the Single Cell Multiome ATAC + Gene Expression assay. Frozen tissue samples were dissociated using a pestle in lysis buffer, and the resulting suspension was passed through a nuclei isolation column. Isolated nuclei were resuspended in debris removal buffer and washed to remove residual debris.

Nuclei viability was assessed by Trypan blue staining and cell counting using a hemocytometer. Nuclei integrity was evaluated under a microscope at 40× or 60× magnification, with viable nuclei characterized by a round morphology and an intact membrane.

Library preparation involved open chromatin transposition and droplet-based gel bead-in-emulsion (GEM) formation, using a Chromium Controller (10X Genomics). Libraries were generated with the Multiome ATAC + Gene Expression kit (10X Genomics), following the manufacturer’s protocol (CG000338, Rev. E). The quality of the resulting ATAC and RNA libraries was assessed using a TapeStation system (Agilent Technologies).

### Sequencing and raw data processing

Multiome libraries were sequenced on the Illumina NovaSeq 6000 platform, following the recommended sequencing depth and read length specifications outlined in the Multiome kit protocol (CG000338). Cell Ranger ARC v2.0.2 (10x Genomics, ‘count’ option with default parameters) was used to filter and align raw reads, identify transposase cut sites, detect accessible chromatin peaks, call cells, and generate raw count matrices for single-cell Multiome ATAC + GEX samples. Alignment was performed using the GRCh38/hg38 reference genome coupled with the Ensembl v98 gene annotation, including reads mapping to intronic regions.

### Quality control

Quality control (QC) was performed using Signac v1.3.0^45^ and Seurat v4.3.0^46^. QC metrics for RNA and Assay for Transposase-Accessible Chromatin (ATAC) modalities were calculated independently but were jointly used to filter cells. A combination of thresholds was established for each sample based on hard cutoffs or on the distribution of each metric within the sample. In the RNA modality, cells were filtered on the number of genes (min = 400, max = mean + 2*sd), unique molecular identifiers (UMIs) (min = 0, max = mean+2*sd), and mitochondrial content (min = 0, max = 20%). In the ATAC modality, cells were filtered on the number of fragments in peaks (min = 400, max = mean+2*sd), transcription start site enrichment (min = mean-2*sd, max = mean+2*sd), and nucleosome signal (min = 0, max = 1.5).

### snRNA-seq data processing

Doublet identification was performed for each sample using scDblFinder v1.8.0^47^. Standard preprocessing and normalization steps were applied individually to each sample using the Seurat v5.1.0^46^ package, including NormalizeData, FindVariableFeatures, ScaleData, RunPCA, RunUMAP, FindNeighbors, and FindClusters. The filtered gene expression matrices were log-normalized with regression for the mitochondrial gene content.

To correct for batch effects across samples, NormalizeData, FindVariableFeatures, and RunPCA were re-applied to the merged dataset. Harmony^48^ v1.2 was used for batch correction and data integration, utilizing default parameters with automatic hyperparameter tuning. Harmony projects cells into a shared embedding space, allowing for clustering based on cell type rather than technical variables such as sequencing batch. Dimensionality reduction was then performed using RunUMAP on the first 40 principal components, as determined by the elbow plot.

Cluster markers were calculated with the FindAllMarkers function, using the Wilcox test on normalized data, with anadjusted p-value < 0.05, average log2 fold-change threshold > 0.25, and a minimum of 50 cells in each group. Differentially expressed genes (DEGs) were obtained using MAST^49^ on normalized data with a percent of cells expressing the gene > 0.1, a minimum of 20 cells in each group, and mitochondrial percentage, brain region, and sequencing batch as latent variables. Significant DEGs exhibited an adjusted p-value < 0.05 and fold-change > 1.5.

### snATAC-seq data processing

Peaks were identified within each sample using MACS2^50^. Standard ATAC-seq preprocessing and normalization were performed using the RunTFIDF, FindTopFeatures, RunSVD, and FindClusters functions from the Signac^45^ v.1.12 package. To account for differences in sequencing depth, the peak count matrix for each sample was normalized using term frequency–inverse document frequency (TF-IDF) via RunTFIDF. Dimensionality reduction was carried out using latent semantic indexing (LSI) with RunSVD.

Prior to data integration, a unified peak set was generated across all samples using MACS2^50^. Peaks in individual samples were quantified using the FeatureMatrix function. Peaks mapping to non-standard chromosomes and ENCODE blacklisted regions were excluded from downstream analyses.

Following this, RunTFIDF, FindTopFeatures, and RunSVD were re-applied to the merged dataset. Harmony was used to integrate the LSI embeddings across samples, using parameters consistent with those applied in RNA integration. UMAP projection was performed using components 2–30, as the first component was found to capture technical variation.

Gene-level chromatin accessibility was computed using the GeneActivity function in Signac^45^ v.1.12.0, which aggregates fragment counts intersecting both gene bodies and promoter regions.

Differentially accessible chromatin regions (DACRs) were computed with FindMarkers using logistic regression, with percent of cells with accessible chromatin > 0.05, a minimum of 20 cells in each group, and the number of peaks and brain region as latent variables. Significant DACRs exhibited adjusted p-value < 0.05 and fold-change > 1.5.

### Cell-type annotation

Cells were annotated at the individual cell level using CoRAL (v3.0.0, https://github.com/fungenomics/CoRAL), a consensus annotation method, using eight single-cell classifier tools: SciBet^45,51^, scClassify^52^, SingleCellNet^53^, Spearman correlation, Support Vector Machines (SVM), ACTINN^54^, scHPL^55^ and scPred^56^. These tools were trained on a reference dataset derived from a human adult single-nucleus atlas^17^ (n = 3,369,219 cells), which includes a nested hierarchy of cell label annotations from the original publication (levels named “SuperClusters” and “Clusters”) and covers several brain regions across the entire adult brain. The training reference was created by excluding groups (author label annotation level “Clusters”) with fewer than 100 cells and downsampling each to a maximum of 150 cells, resulting in a reference of n = 67,696 cells and n= 453 labels.

A consensus label for each cell was computed using a modified version of the CAWPE^57^ algorithm as follows: first, the F1 accuracy measure was computed for each single-cell classifier by performing a 5-fold cross-validation in the reference data set and calculating the average F1 across folds. Second, each classifier was trained on the reference data set and used to predict reference labels with a confidence score (Sl), as follows:

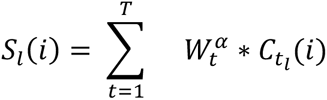

Where *i* is an individual cell, W_t_ is the cross-validation accuracy (F1) measure for tool *t*, α is a hyper parameter (set to 4), and 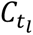 is the predicted confidence score of being of label *l* assigned by the tool *t*. The scores were then grouped and averaged into the SuperClusters level annotation based on the authors’ pre-defined cell-type hierarchy. The final label for each cell was assigned to the SuperClusters category with the highest score.

### Multimodal data integration

To integrate gene expression and chromatin accessibility data, we employed Weighted Nearest Neighbor (WNN) analysis, an unsupervised method that assesses the relative contribution of each modality and constructs a unified neighbor graph representing both RNA and ATAC data. WNN analysis was performed using the FindMultiModalNeighbors function with 20 neighbors (k.nn), based on UMAP embeddings obtained following Harmony^4^integration. The resulting WNN graph was used to generate a joint UMAP visualization. Clustering was then performed using the FindClusters function with the smart local moving (SLM) algorithm, with a resolution of 2.

### Post-clustering Quality Control

Cells classified as doublets, as well as clusters exhibiting a high proportion of doublet cells, were excluded from further analysis. Clusters containing fewer than 50 nuclei or showing significant enrichment for mitochondrial markers were also removed. Additionally, cell types represented by fewer than 50 cells or identified in only a single patient were excluded. Cells annotated as ‘miscellaneous’ or ‘splatter’, reflecting non-specific cell types present in the reference dataset, were removed. Following manual inspection of canonical markers, cells annotated as ‘Bergmann glia’ were reclassified as astrocytes. Furthermore, region-specific cell types—’Amygdala excitatory’ in the amygdala, ‘Upper-layer IT’, ‘Deep-layer IT’, ‘Deep-layer CT/6b’, and ‘Deep-layer NP’ in the cortex, as well as ‘Hippocampal CA1-3’ and ‘Hippocampal DG’ in the hippocampus—that were detected outside of their respective regions were also removed.

### Cellular composition analysis

Cell type/subcluster counts were quantified across brain region and age/period using the R package sccomp^58^ testing for donor age, brain region, sequencing batch, and sex. Briefly, sccomp^58^ models counts while accounting for group-specific variability by employing a Bayesian model based on sum-constrained beta-binomial distributions.

### Gene regulatory networks construction

We used SCENIC+^59^ v.1.0a2 with Python v.3.11.8 for enhancer-driven gene regulatory network construction. To reduce computational cost, we randomly sampled 20,000 quality-controlled nuclei for this analysis. Samples were preprocessed with pycisTopic v.2.0.2a. Cells were not filtered at this stage, since QC was previously performed (as described in the section “snRNA-seq analysis” in Methods). We generated pseudobulk profiles for each cell type and used MACS2 to call peaks, from which blacklist regions were removed, resulting in 899,049 peaks. Parallel Latent Dirichlet Allocation (LDA) with Machine Learning for Language Toolkit (MALLET) was used for topic modeling with the function run_cgs_models_mallet, using 200 iterations and a symmetric Dirichlet hyperparameter for topic proportions (alpha) of 50. Models were evaluated with evaluate_models and 30 was chosen as the optimal number of topics. Dimensionality reduction was performed using UMAP to visualize cell–topic contributions to the model, followed by Leiden clustering with a resolution parameter of 1.2. Topic binarization was conducted using the Otsu thresholding method and by selecting the top 3,000 most contributing regions. Region accessibility was imputed and normalized using scale factors of 10⁶ and 10⁴, respectively. Differentially accessible regions (DARs) were identified with the find_diff_features function applied to highly variable regions (minimum dispersion = 0.05; mean accessibility between 0.0125 and 3), using an adjusted *p*-value < 0.05 and fold change > 1.5 as significance thresholds.

Single-nucleus RNA-seq (snRNA-seq) data were processed using the Scanpy v1.8.2 framework^60^. Data were normalized with normalize_total (target sum = 10⁴) followed by log-transformation with log1p. Highly variable genes were selected using highly_variable_genes with a mean expression between 0.0125 and 3 and minimum dispersion of 0.5. Data scaling was performed with the scale function (maximum value = 10). Dimensionality reduction was carried out with the pca and umap functions using default parameters, and graphs were constructed with the neighbors function (n_neighbors = 20; n_pcs = 40).

Custom cisTarget ranking and scoring databases were generated using consensus peak regions. Cluster-Buster^61^ was employed to score genomic regions against the Aerts lab motif collection (v10nr_clust_public; https://resources.aertslab.org/cistarget/motif_collections/v10nr_clust_public/v10nr_clust_public.zip). FASTA sequences were generated from consensus regions padded by 1 kb of background sequence.

The processed pyCisTopic and snRNA-seq objects, along with region sets, cisTarget databases, and motif annotations, were provided as input to SCENIC+^59^, which was executed as a Snakemake pipeline (seed = 1234) with default parameters for data preparation, motif enrichment, and parameter inference. This analysis yielded 477 eRegulons, defined as transcription factors (TFs) and their associated genomic regions and target genes. SCENIC+ metadata for these eRegulons can be found in Supplementary Data 1.

Dimensionality reduction of eRegulon activity was performed by applying neighbors and umap (default parameters) to the matrix of gene-based eRegulon activity scores per cell. To identify cell type– and brain region–specific eRegulons, Regulon Specificity Scores (RSS) were computed using cell type and brain region as grouping variables. For network visualization, the top 200 highest-ranked triplets (TF–region–gene combinations) among the 20 eRegulons with the highest RSS values were imported into Cytoscape^62^ v3.10.3.

### Identification of age-associated eRegulons

Age-associated eRegulons were identified using psupertime^63^ v0.2.6. Briefly, psupertime fits an ordered logistic regression model to identify features (e.g., genes or cis-regulatory elements) that recapitulate the group-level temporal ordering of cells. Gene- and region-based eRegulon activity score (AUC) matrices, together with biological age, were provided as input to psupertime. The algorithm estimates pseudotime values per cell and reports eRegulons with significant non-zero age coefficients. To identify eRegulons displaying an approximately linear relationship with age, linear models were fit between a-scored activity scores (AUC) and pseudotime across cells, and significant eRegulons with R² > 0.1 were retained. Age-associated eRegulons were defined as activators exhibiting a mean activity score > 0.1, mean TF expression > 0.1, and TF expression detected in >25% of cells. For network visualization, we selected the top 200 highest-ranked triplets (TF–region–gene combinations) among the age-associated eRegulons of each cell type.

### Cis-regulatory elements overlap analysis

The overlap between expression quantitative trait loci (eQTLs) and genetic risk variants was evaluated with R packages GenomicRanges^64^ v.1.46.1 and regioneR^65^ v.1.46.1. When necessary, genomic coordinates were converted from hg38 to hg19 using the liftOver function of the R package rtracklayer^66^ v.1.54.0 and the chain file hg38ToHg19.over.chain. Briefly, regioneR performs permutation-based statistical testing to assess whether the observed overlap between two sets of genomic regions exceeds that expected by chance. The permTest function was applied using 10,000 permutations. For cis-eQTLs, the parameter per_chromosome was set to TRUE, ensuring that randomizations were constrained within each chromosome. eQTLs were obtained from the MetaBrain project^67^ and genetic risk variants were obtained from previous studies^68–77^.

## Author contributions

I.C.G., T.V.W., B.C., S.H., and D.F.T.V. performed single-cell data analyses. T.C. de O., L.H.B. dos S. performed ancestry analysis. M.L. performed experimental single-nucleus assays. C.L.Y., A.C.A., E.G., H.T., M.K.M.A., F.C., and I.L-C contributed to clinical sample collection. F.R. performed histopathological analyses. I.C.G. and D.F.T.V. wrote the manuscript, with input from all authors. D.F.T.V. acquired funding and designed the study. All authors reviewed and approved the final version of the manuscript.

## SUPPLEMENTARY FIGURES

**Figure S1.**
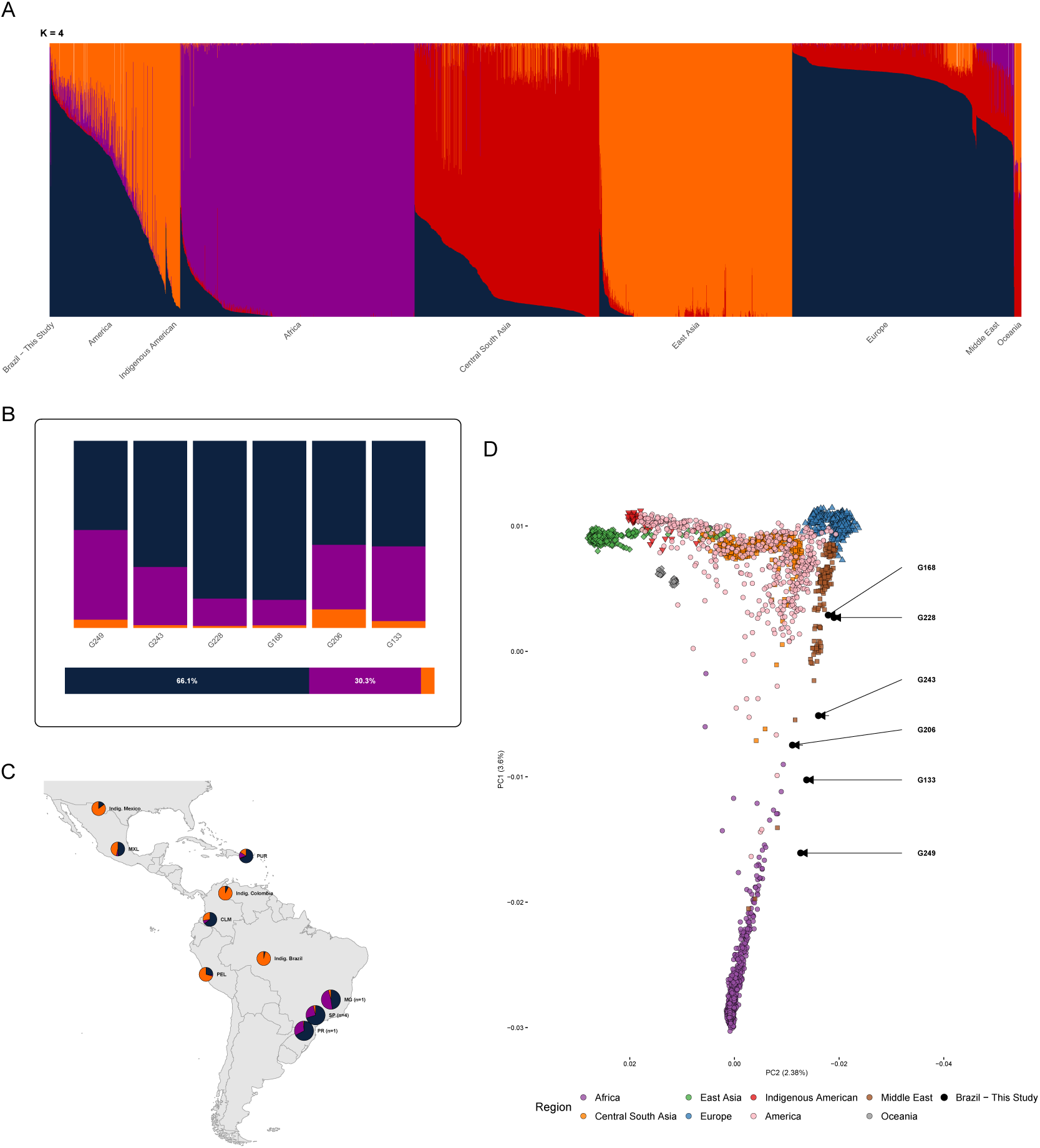
Genetic ancestry of Brazilian donors sequenced in this study, in the context of global reference populations. **(A)** ADMIXTURE barplot showing estimated ancestry proportions at K = 4. Unsupervised clustering was performed on the LD-pruned autosomal SNP dataset comprising the six study donors merged with the HGDP+1KGP reference panel (n = 4,157). Each vertical bar represents an individual, with colored segments indicating the proportional contribution of three inferred ancestral components. Individuals are grouped by population. **(B)** ADMIXTURE barplot of the six Brazilian donors included in this study, with mean proportions shown in the horizontal bar below. Brazilian donors have an admixed profile with predominantly European ancestry, followed by African and Indigenous American contributions. **(C)** Geographic distribution of mean ancestry proportions across the Americas, displaying pie charts for Brazilian donors by state (this study; states include Minas Gerais (MG), Paraná (PR), and São Paulo (SP)), Indigenous American populations (Maya, Pima, Karitiana, Surui, Colombian), and admixed Latin American populations from the HGDP+1KGP. **(D)** PCA performed using SNPRelate showing the distribution of the six Brazilian donors from this study relative to global reference populations from the HGDP+1KGP panel. Arrows with donor IDs indicate the samples sequenced in this study. Human Genome Diversity Project, HGDP; 1000 Genomes Project (1KGP)

**Figure S2.**
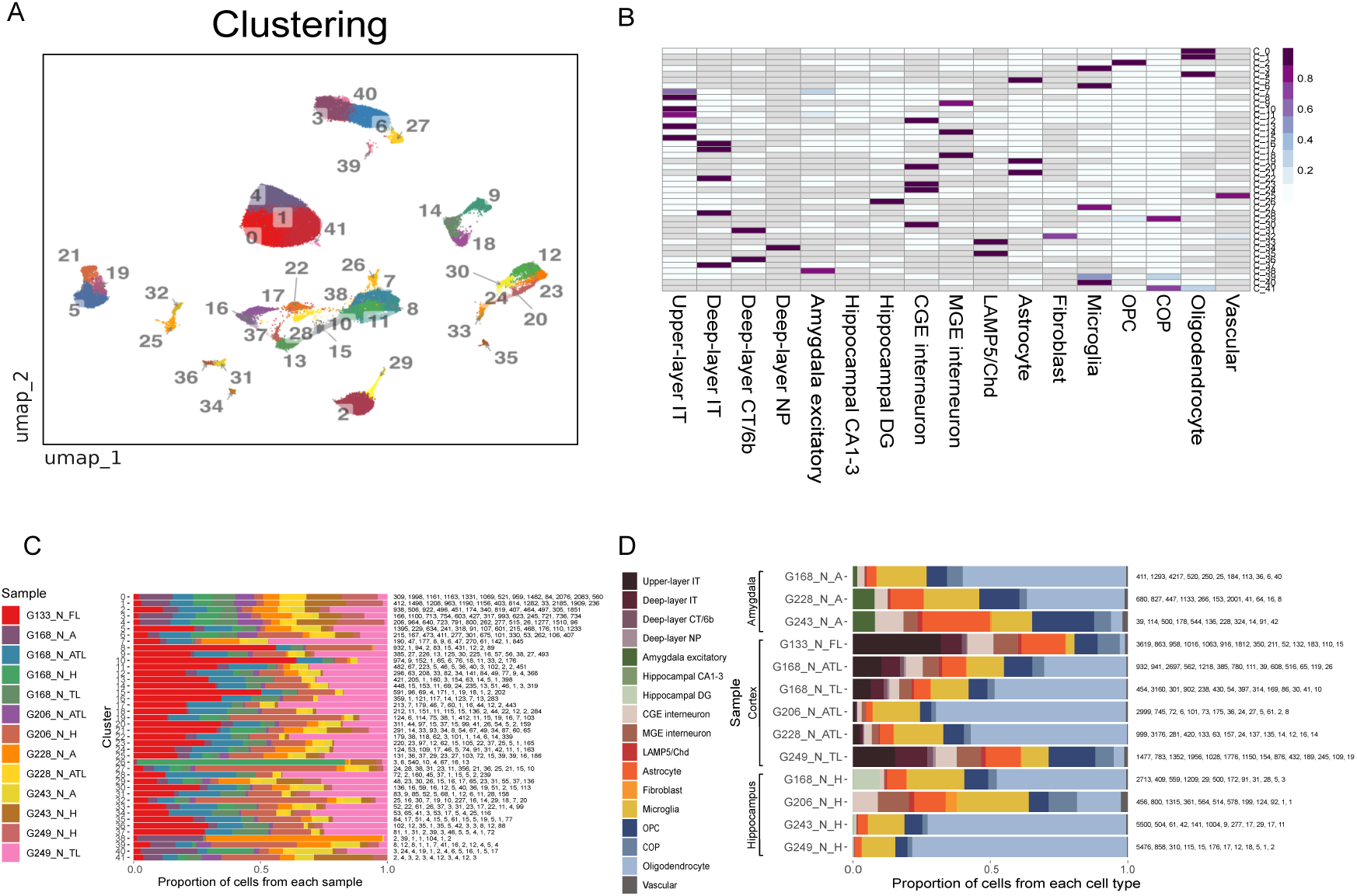
Clustering and annotation. (A) Uniform Manifold Approximation and Projection (UMAP) visualization of nuclei clusters obtained from multimodal integration. Clustering was performed using a joint graph derived from Weighted Nearest Neighbor (WNN) analysis combining RNA and ATAC modalities. (B) Heatmap showing final cluster annotations, where color represents the proportion of each cell type within a given cluster. (C) Bar plots illustrating the distribution of donor samples across clusters. (D) Bar plots showing the relative proportions of cell types within biological samples, grouped by brain region.

**Figure S3.**
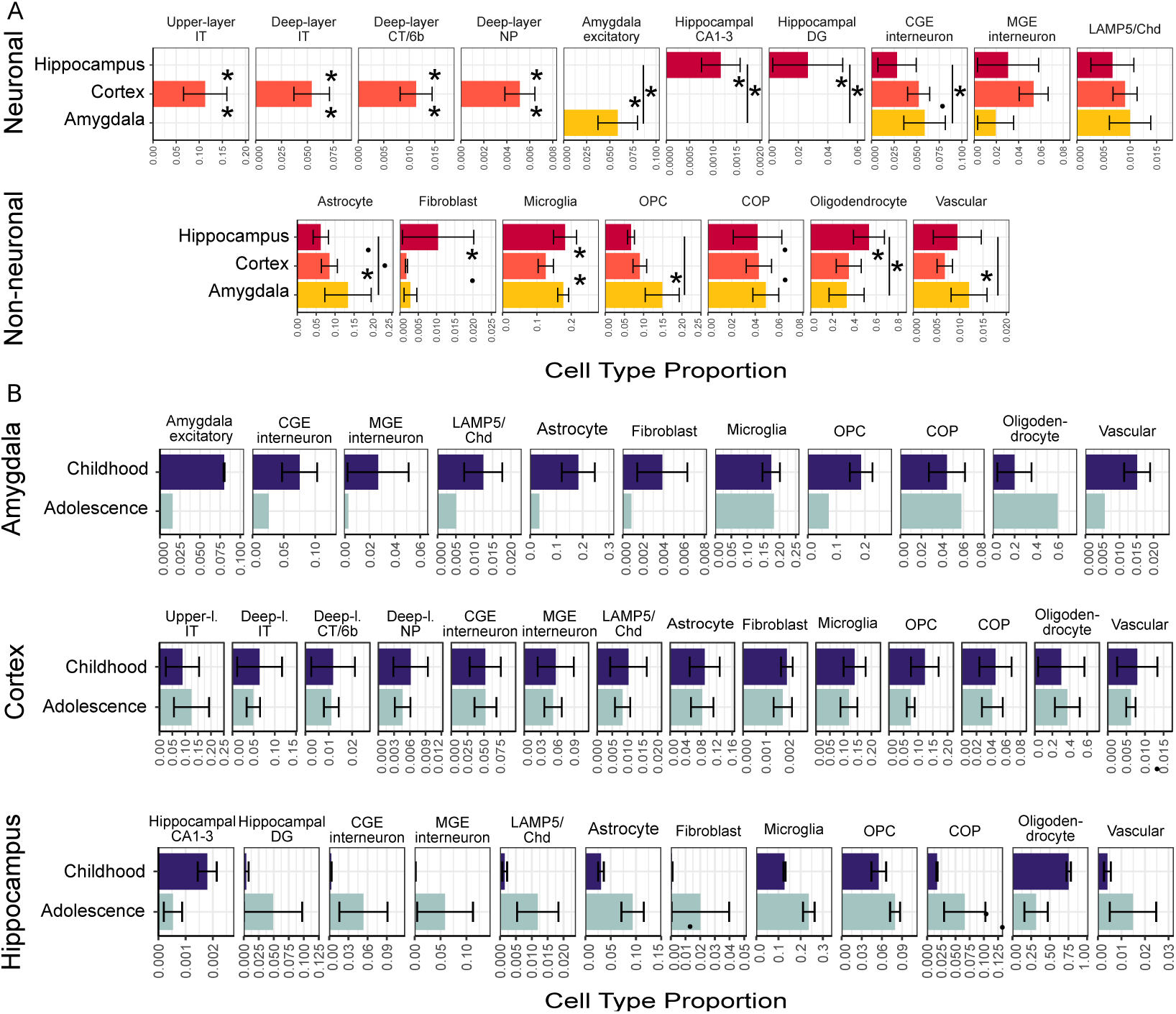
Cellular composition of brain regions during the transition from childhood to adolescence. (A) Bar plots showing the relative abundance of major cell types in childhood (≤10 years) and adolescence (> 10 years) across the amygdala (top), cortex (middle), and hippocampus (bottom). Panel titles indicate the cell type. The *x*-axis represents the proportion of that cell type, while the *y*-axis and bar color denote the developmental period. Error bars mark standard error of the mean and asterisks/dots mark significant differences in abundance across donor age (* p < 0.05; ׄ p < 0.1). (B) Bar plots illustrating cell type composition across brain regions. Panel titles indicate the cell type. The *x*-axis shows the proportion of each cell type, and the *y*-axis and bar color correspond to the brain region. Error bars mark standard error of the mean and asterisks/dots mark significant differences in abundance across brain regions (* p < 0.05; ׄ p < 0.1).

**Figure S4.**
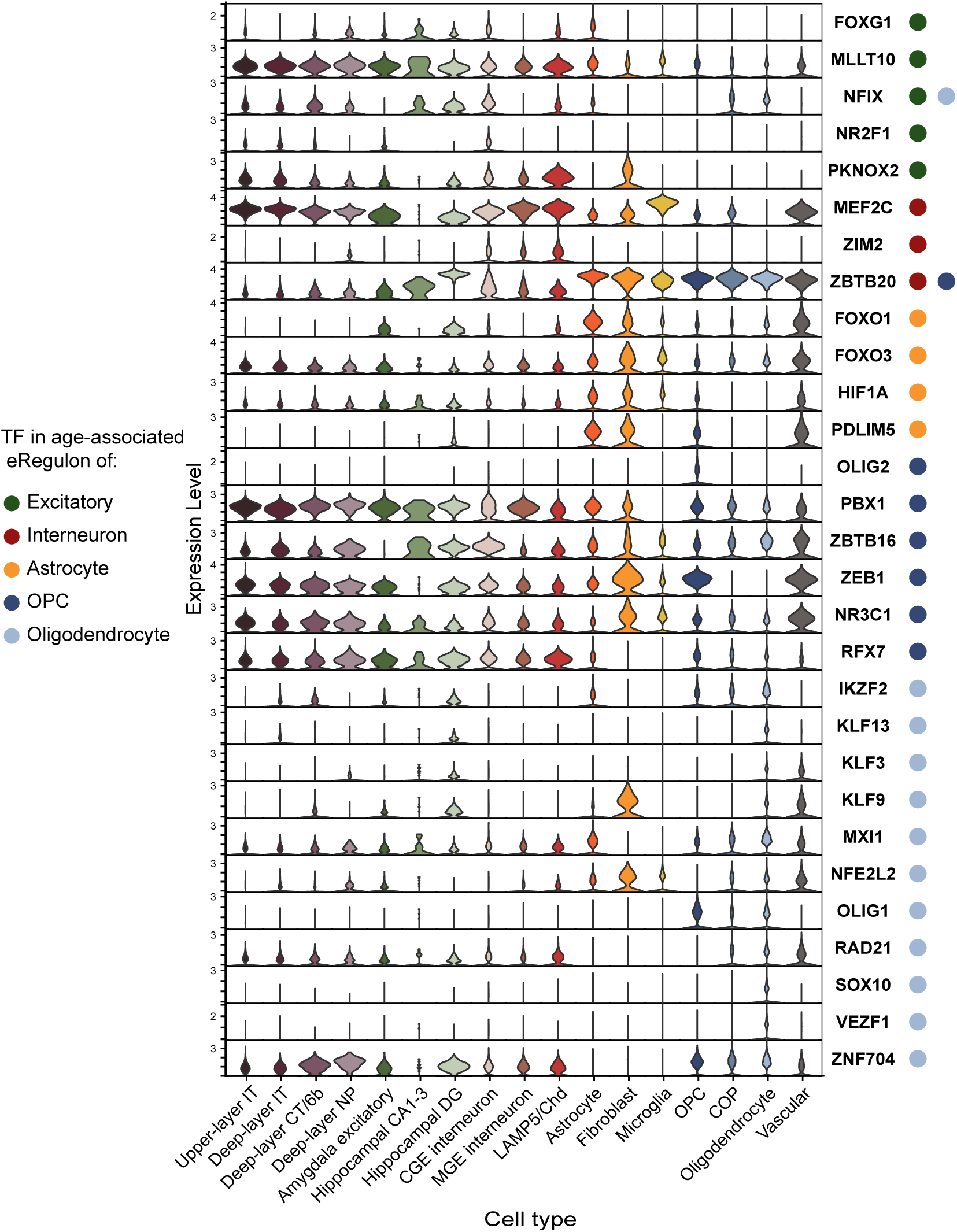
Expression of transcription factors driving adolescence-associated eRegulons. Violin plots showing the expression levels of transcription factors (TFs) in adolescence-associated eRegulons across brain cell types. The *x*-axis and plot colors indicate cell identity, and the *y*-axis shows normalized TF expression. TF names are listed on the right. Colored dots adjacent to each TF label denote the cell type(s) in which the eRegulon was identified.

## SUPPLEMENTARY DATA

**Supplementary Data 1.** SCENIC+ metadata for all eRegulons.

## SUPPLEMENTARY TABLES

**Supplementary Table 1.** Age-associated eRegulons statistics.

**Supplementary Table 2.** Age-associated eRegulons SCENIC+ metadata.

**Supplementary Table 3.** eQTLs overlapping age-associated target regions.

**Supplementary Table 4.** Gene Ontology terms enriched in age-associated target genes per cell type and eRegulon.

**Supplementary Table 5.** Gene sets containing genetic risk variants.

**Supplementary Table 6.** Genetic risk variants overlapping age-associated target regions.

## REFERENCES

1. Dong, H.-M., Margulies, D. S., Zuo, X.-N. & Holmes, A. J. Shifting gradients of macroscale cortical organization mark the transition from childhood to adolescence. Proceedings of the National Academy of Sciences 118, e2024448118 (2021).

2. López-Vicente, M. et al. Developmental Changes in Dynamic Functional Connectivity From Childhood Into Adolescence. Front. Syst. Neurosci. 15, (2021).

3. Neniskyte, U. & Gross, C. T. Errant gardeners: glial-cell-dependent synaptic pruning and neurodevelopmental disorders. Nat Rev Neurosci 18, 658–670 (2017).

4. Faust, T. E., Gunner, G. & Schafer, D. P. Mechanisms governing activity-dependent synaptic pruning in the developing mammalian CNS. Nat Rev Neurosci 22, 657–673 (2021).

5. Akinlaja, Y. O. & Nishiyama, A. Glial modulation of synapse development and plasticity: oligodendrocyte precursor cells as a new player in the synaptic quintet. Front. Cell Dev. Biol. 12, (2024).

6. de Faria, O. et al. Periods of synchronized myelin changes shape brain function and plasticity. Nat Neurosci 24, 1508–1521 (2021).

7. Meyer, H. C. & Lee, F. S. Translating Developmental Neuroscience to Understand Risk for Psychiatric Disorders. American Journal of Psychiatry 180, 540–547 (2023).

8. Howes, O. D. & Onwordi, E. C. The synaptic hypothesis of schizophrenia version III: a master mechanism. Mol Psychiatry 28, 1843–1856 (2023).

9. Polioudakis, D. et al. A Single-Cell Transcriptomic Atlas of Human Neocortical Development during Mid-gestation. Neuron 103, 785–801.e8 (2019).

10. Domcke, S. et al. A human cell atlas of fetal chromatin accessibility. Science 370, eaba7612 (2020).

11. Eze, U. C., Bhaduri, A., Haeussler, M., Nowakowski, T. J. & Kriegstein, A. R. Single-cell atlas of early human brain development highlights heterogeneity of human neuroepithelial cells and early radial glia. Nat Neurosci 24, 584–594 (2021).

12. Ramos, S. I. et al. An atlas of late prenatal human neurodevelopment resolved by single-nucleus transcriptomics. Nat Commun 13, 7671 (2022).

13. Bruggen, D. van et al. Developmental landscape of human forebrain at a single-cell level identifies early waves of oligodendrogenesis. Developmental Cell 57, 1421–1436.e5 (2022).

14. Zeng, B. et al. The single-cell and spatial transcriptional landscape of human gastrulation and early brain development. Cell Stem Cell 30, 851–866.e7 (2023).

15. Mannens, C. C. A. et al. Chromatin accessibility during human first-trimester neurodevelopment. Nature 647, 179–186 (2025).

16. Nunes, K. et al. Admixture’s impact on Brazilian population evolution and health. Science 388, eadl3564 (2025).

17. Siletti, K. et al. Transcriptomic diversity of cell types across the adult human brain. Science 382, eadd7046 (2023).

18. Chen, J. et al. Single-nucleus transcriptome atlas of the basal forebrain reveals diverse ageing-related pathways. Brain 148, 2493–2508 (2025).

19. Li, P., Goodwin, A., Halushka, P. & Fan, H. Unveiling the role of Fli-1 in controlling astrocyte dysfunction and transformation in Alzheimer’s disease. Alzheimer’s & Dementia 20, e086131 (2024).

20. Wang, S., Pan, L., Sun, C., Ma, C. & Pan, H. Balancing Microglial Density and Activation in Central Nervous System Development and Disease. Current Issues in Molecular Biology 47, 344 (2025).

21. Zhu, Z., Meng, M., Mo, S., Wang, X. & Qiao, L. M2 Microglia-Derived Exosomal miR-144-5p Attenuates White Matter Injury in Preterm Infants by Regulating the PTEN/AKT Pathway Through KLF12. Mol Biotechnol 10.1007/s12033-025-01364-1 (2025) doi:10.1007/s12033-025-01364-1.

22. Weider, M. & Wegner, M. SoxE factors: Transcriptional regulators of neural differentiation and nervous system development. Seminars in Cell & Developmental Biology 63, 35–42 (2017).

23. Bernhardt, C. et al. KLF9 and KLF13 transcription factors boost myelin gene expression in oligodendrocytes as partners of SOX10 and MYRF. Nucleic Acids Res 50, 11509–11528 (2022).

24. Du, S. et al. FoxO3 deficiency in cortical astrocytes leads to impaired lipid metabolism and aggravated amyloid pathology. Aging Cell 20, e13432 (2021).

25. Doan, K. V. et al. Astrocytic FoxO1 in the hypothalamus regulates metabolic homeostasis by coordinating neuropeptide Y neuron activity. Glia 71, 2735–2752 (2023).

26. Du, J. et al. S100B is selectively expressed by gray matter protoplasmic astrocytes and myelinating oligodendrocytes in the developing CNS. Mol Brain 14, 154 (2021).

27. Deloulme, J. C. et al. Nuclear expression of S100B in oligodendrocyte progenitor cells correlates with differentiation toward the oligodendroglial lineage and modulates oligodendrocytes maturation. Molecular and Cellular Neuroscience 27, 453–465 (2004).

28. Herring, C. A. et al. Human prefrontal cortex gene regulatory dynamics from gestation to adulthood at single-cell resolution. Cell 185, 4428–4447.e28 (2022).

29. Wang, L. et al. Molecular and cellular dynamics of the developing human neocortex. Nature 647, 169–178 (2025).

30. Mesman, S. & Smidt, M. P. Tcf12 Is Involved in Early Cell-Fate Determination and Subset Specification of Midbrain Dopamine Neurons. Front. Mol. Neurosci. 10, (2017).

31. Cattane, N. et al. Altered Gene Expression in Schizophrenia: Findings from Transcriptional Signatures in Fibroblasts and Blood. PLOS ONE 10, e0116686 (2015).

32. Sock, E. & Wegner, M. Using the lineage determinants Olig2 and Sox10 to explore transcriptional regulation of oligodendrocyte development. Developmental Neurobiology 81, 892–901 (2021).

33. Turnescu, T. et al. Sox8 and Sox10 jointly maintain myelin gene expression in oligodendrocytes. Glia 66, 279–294 (2018).

34. Yu, M., et al. Integrative multi-omic profiling of adult mouse brain endothelial cells and potential implications in Alzheimer’s disease. Cell Reports 42, (2023).

35. Harris, H. K. et al. Disruption of RFX family transcription factors causes autism, attention-deficit/hyperactivity disorder, intellectual disability, and dysregulated behavior. Genet Med 23, 1028–1040 (2021).

36. Liu, A. et al. Single-nucleus multiomics reveals the disrupted regulatory programs in three brain regions of sporadic early-onset Alzheimer’s disease. bioRxiv 2024.06.25.600720 (2024) doi:10.1101/2024.06.25.600720.

37. Clarence, T. et al. Multiomic single-cell profiling identifies critical regulators of postnatal brain. Nat Genet 57, 591–603 (2025).

38. Hong, S., Dissing-Olesen, L. & Stevens, B. New insights on the role of microglia in synaptic pruning in health and disease. Current Opinion in Neurobiology 36, 128–134 (2016).

39. Lemke, J. R. et al. GRIN2A null variants confer a high risk for early-onset schizophrenia and other mental disorders and potentially enable precision therapy. Mol Psychiatry 31, 374–382 (2026).

40. Koenig, Z. et al. A harmonized public resource of deeply sequenced diverse human genomes. Genome Res. 34, 796–809 (2024).

41. Chang, C. C. et al. Second-generation PLINK: rising to the challenge of larger and richer datasets. Gigascience 4, 7 (2015).

42. Purcell, S. et al. PLINK: A Tool Set for Whole-Genome Association and Population-Based Linkage Analyses. The American Journal of Human Genetics 81, 559–575 (2007).

43. Zheng, X. et al. A high-performance computing toolset for relatedness and principal component analysis of SNP data. Bioinformatics 28, 3326–3328 (2012).

44. Alexander, D. H., Novembre, J. & Lange, K. Fast model-based estimation of ancestry in unrelated individuals. Genome Res 19, 1655–1664 (2009).

45. Stuart, T., Srivastava, A., Madad, S., Lareau, C. A. & Satija, R. Single-cell chromatin state analysis with Signac. Nat Methods 18, 1333–1341 (2021).

46. Hao, Y. et al. Integrated analysis of multimodal single-cell data. Cell 184, 3573–3587.e29 (2021).

47. Germain, P.-L., Lun, A., Garcia Meixide, C., Macnair, W. & Robinson, M. D. Doublet identification in single-cell sequencing data using scDblFinder. F1000Res 10, 979 (2021).

48. Korsunsky, I. et al. Fast, sensitive and accurate integration of single-cell data with Harmony. Nat Methods 16, 1289–1296 (2019).

49. Finak, G. et al. MAST: a flexible statistical framework for assessing transcriptional changes and characterizing heterogeneity in single-cell RNA sequencing data. Genome Biology 16, 278 (2015).

50. Zhang, Y. et al. Model-based Analysis of ChIP-Seq (MACS). Genome Biology 9, R137 (2008).

51. Li, C. et al. SciBet as a portable and fast single cell type identifier. Nat Commun 11, 1818 (2020).

52. Lin, Y. et al. scClassify: sample size estimation and multiscale classification of cells using single and multiple reference. Mol Syst Biol 16, e9389 (2020).

53. Tan, Y. & Cahan, P. SingleCellNet: A Computational Tool to Classify Single Cell RNA-Seq Data Across Platforms and Across Species. Cell Syst 9, 207–213.e2 (2019).

54. Ma, F. & Pellegrini, M. ACTINN: automated identification of cell types in single cell RNA sequencing. Bioinformatics 36, 533–538 (2020).

55. Michielsen, L., Reinders, M. J. T. & Mahfouz, A. Hierarchical progressive learning of cell identities in single-cell data. Nat Commun 12, 2799 (2021).

56. Alquicira-Hernandez, J., Sathe, A., Ji, H. P., Nguyen, Q. & Powell, J. E. scPred: accurate supervised method for cell-type classification from single-cell RNA-seq data. Genome Biology 20, 264 (2019).

57. Large, J., Lines, J. & Bagnall, A. A probabilistic classifier ensemble weighting scheme based on cross-validated accuracy estimates. Data Min Knowl Disc 33, 1674–1709 (2019).

58. Mangiola, S. et al. sccomp: Robust differential composition and variability analysis for single-cell data. Proceedings of the National Academy of Sciences 120, e2203828120 (2023).

59. Bravo González-Blas, C., et al. SCENIC+: single-cell multiomic inference of enhancers and gene regulatory networks. Nat Methods 20, 1355–1367 (2023).

60. Wolf, F. A., Angerer, P. & Theis, F. J. SCANPY: large-scale single-cell gene expression data analysis. Genome Biol 19, 15 (2018).

61. Frith, M. C., Li, M. C. & Weng, Z. Cluster-Buster: finding dense clusters of motifs in DNA sequences. Nucleic Acids Res 31, 3666–3668 (2003).

62. Shannon, P. et al. Cytoscape: A Software Environment for Integrated Models of Biomolecular Interaction Networks. Genome Res. 13, 2498–2504 (2003).

63. Macnair, W., Gupta, R. & Claassen, M. psupertime: supervised pseudotime analysis for time-series single-cell RNA-seq data. Bioinformatics 38, i290–i298 (2022).

64. Lawrence, M. et al. Software for Computing and Annotating Genomic Ranges. PLOS Computational Biology 9, e1003118 (2013).

65. Gel, B. et al. regioneR: an R/Bioconductor package for the association analysis of genomic regions based on permutation tests. Bioinformatics 32, 289–291 (2016).

66. Lawrence, M., Gentleman, R. & Carey, V. rtracklayer: an R package for interfacing with genome browsers. Bioinformatics 25, 1841–1842 (2009).

67. de Klein, N. et al. Brain expression quantitative trait locus and network analyses reveal downstream effects and putative drivers for brain-related diseases. Nat Genet 55, 377–388 (2023).

68. Corces, M. R. et al. Single-cell epigenomic analyses implicate candidate causal variants at inherited risk loci for Alzheimer’s and Parkinson’s diseases. Nat Genet 52, 1158–1168 (2020).

69. Chawla, A. et al. Single-nucleus chromatin accessibility profiling identifies cell types and functional variants contributing to major depression. Nat Genet 57, 1890–1904 (2025).

70. Grove, J. et al. Identification of common genetic risk variants for autism spectrum disorder. Nat Genet 51, 431–444 (2019).

71. O’Connell, K. S. et al. Genomics yields biological and phenotypic insights into bipolar disorder. Nature 639, 968–975 (2025).

72. Demontis, D. et al. Genome-wide analyses of ADHD identify 27 risk loci, refine the genetic architecture and implicate several cognitive domains. Nat Genet 55, 198–208 (2023).

73. Stevelink, R. et al. GWAS meta-analysis of over 29,000 people with epilepsy identifies 26 risk loci and subtype-specific genetic architecture. Nat Genet 55, 1471–1482 (2023).

74. Savage, J. E. et al. Genome-wide association meta-analysis in 269,867 individuals identifies new genetic and functional links to intelligence. Nat Genet 50, 912–919 (2018).

75. Pardiñas, A. F. et al. Common schizophrenia alleles are enriched in mutation-intolerant genes and in regions under strong background selection. Nat Genet 50, 381–389 (2018).

76. Nagel, M. et al. Meta-analysis of genome-wide association studies for neuroticism in 449,484 individuals identifies novel genetic loci and pathways. Nat Genet 50, 920–927 (2018).

77. Posthuma, D. & International Obsessive Compulsive Disorder Foundation Genetics Collaborative (IOCDF-GC) and OCD Collaborative Genetics Association Studies (OCGAS). Revealing the complex genetic architecture of obsessive-compulsive disorder using meta-analysis. Molecular Psychiatry 23, 1181–1188 (2018).

